# Assessing the impact of autologous neutralizing antibodies on rebound dynamics in postnatally SHIV-infected ART-treated infant rhesus macaques

**DOI:** 10.1101/2024.06.01.596971

**Authors:** Ellie Mainou, Stella J Berendam, Veronica Obregon-Perko, Emilie A Uffman, Caroline T Phan, George M Shaw, Katharine J Bar, Mithra R Kumar, Emily J Fray, Janet M Siliciano, Robert F Siliciano, Guido Silvestri, Sallie R Permar, Genevieve G Fouda, Janice McCarthy, Ann Chahroudi, Cliburn Chan, Jessica M Conway

**Affiliations:** Department of Biology, Pennsylvania State University, University Park, PA, USA; GlaxoKlineSmith, Rockville, MD, USA; FlowJo, Altanta, GA, USA; Duke Human Vaccine Institute, Duke University Medical Center, Durham, NC, USA; Perelman School of Medicine, University of Pennsylvania, Philadelphia, Pennsylvania, USA; Department of Pharmacology and Molecular Sciences, Johns Hopkins University School of Medicine, Baltimore, MD, USA; Department of Biochemistry and Molecular Biology, Johns Hopkins University School of Medicine, Baltimore, MD, USA; Department of Medicine, Johns Hopkins University School of Medicine, Baltimore, MD, USA; Yerkes National Primate Research Center, Emory University, Atlanta, Georgia, USA; Department of Pediatrics, Weill Cornell Medicine, New York, NY, USA; Department of Biostatistics and Bioinformatics, Duke University Medical Center, Durham, NC, USA; Department of Pediatrics, Emory University School of Medicine, Atlanta, GA 30322, USA; Department of Mathematics, Pennsylvania State University, University Park, PA, USA

## Abstract

The presence of antibodies against HIV in infected children is associated with a greater capacity to control viremia in the absence of therapy. While the benefits of early antiretroviral treatment (ART) in infants are well documented, early ART may interfere with the development of antibody responses. In contrast to adults, early treated children lack detectable HIV-specific antibodies, suggesting a fundamental difference in HIV pathogenesis. Despite this potential adverse effect, early ART may decrease the size of the latent reservoir established early in infection in infants, which can be beneficial in viral control. Understanding the virologic and immunologic aspects of pediatric HIV is crucial to inform innovative targeted strategies for treating children living with HIV. In this study, we investigate how ART initiation time sets the stage for trade-offs in the latent reservoir establishment and the development of humoral immunity and how these, in turn, affect post-treatment dynamics. We also elucidate the biological function of antibodies in pediatric HIV. We employ mathematical modeling coupled with experimental data from an infant nonhuman primate Simian/Human Immunodeficiency Virus (SHIV) infection model. Infant Rhesus macaques (RMs) were orally challenged with SHIV.C.CH505 375H dCT four weeks after birth and started treatment at different times after infection. In addition to viral load measurements, antibody responses and latent reservoir sizes were measured. We estimate model parameters by fitting viral load measurements to the standard HIV viral dynamics model within a nonlinear fixed effects framework. This approach allows us to capture differences between rhesus macaques (RMs) that develop antibody responses or exhibit high latent reservoir sizes compared to those that do not. We find that neutralizing antibody responses are associated with increased viral clearance and decreased viral infectivity but decreased death rate of infected cells. In addition, the presence of detectable latent reservoir is associated with less robust immune responses. These results demonstrate that both immune response and latent reservoir dynamics are needed to understand post-rebound dynamics and point to the necessity of a comprehensive approach in tailoring personalized medical interventions.

## 1 Introduction

HIV-1 continues to pose a significant global health challenge, affecting 38.4 million individuals as of 2021, according to the World Health Organization [1]. Among these, 1.7 million are children, as reported by UNAIDS [2].

The progression of HIV infection and the way infants respond to treatment vary significantly from that of adults. In adults, HIV infection is a slow process with a median time to progression to AIDS of 10 years. Plasma viral load decreases 100-1000-fold post-peak viremia and remains relatively stable at the setpoint for several years [3, 4]. In contrast, infant plasma viremia increases to levels that are much higher than adults [5, 6] and declines very slowly reaching a setpoint at around 5 years of age [7, 8]. In addition, rates of clinical disease progression also differ from adults. For example, 20 *−* 30% of children experienced rapid progression to AIDS or death in the first year of life [7]. These rates of diseases progression also tend to vary depending on the age of the child: a 1-year with 10% CD4+ T cells has a 40% risk of progression to AIDS within in year compared to a 10-year old with same laboratory values who has a 7.4% risk for developing AIDS within one year [7, 9]. These notable distinctions are mostly attributed to differences in the immunologic ontology of infants as well as the rapid CD4+ T cell expansion that accompanies somatic growth [7]. As part of this study, we aim to elucidate the effect of antibodies in pediatric HIV after treatment interruption. In adults, neutralizing antibodies to HIV have been reported to develop 2-4 years after virus transmission [10]. Although only a small fraction of HIV-1 infected individuals develops broad and potent neutralization response, the overall neutralizing response is a continuum with most individuals presenting some level of neutralizing antibodies [11]. In contrast, recent studies in infants demonstrated their ability to develop broadly neutralizing antibodies within the first 24 months of infection, suggesting that neonatal B cells and T helper cells are equipped to respond to HIV-1 envelope, an immune response that can be harnessed in strategies towards a cure [12]. Crucially, the presence of antibodies capable of mediating antibody-dependent cellular cytotoxicity (ADCC) is associated with a better disease outcome in HIV-infected infants, despite the delayed development of such antibodies [13–15].

Antiretroviral treatment has important effects on host-virus dynamics that manifest following treatment interruption. ART is not a cure, and treatment interruption leads to rebound of viremia to levels typical of chronic infection, with remarkable heterogeneity in rebound times [16]. According to some data in HIV infected children treated soon after birth, early ART may also delay viral rebound after treatment interruption also in infants. For instance, the “Mississippi baby” was treated 30 hours after birth and discontinued at 18 months of age, and no detectable viremia was observed for 28 months after treatment discontinuation [17, 18]. This case study sparked hopes for functional cure of pediatric HIV, i.e. sustained suppression of viremia without ART, through very early administration of ART. Additional reports of potential pediatric HIV-1 remission, through early ART followed [19, 20], but were not attributed to a specific mechanism (cellular or humoral immunity). Despite such promising cases [20, 21], a generalizable approach for a functional cure in infants and children has not been achieved [22–24]. In contrast to adults, early-treated children lack detectable HIV-specific antibodies (and CD8+ T cell responses), suggesting a fundamental difference in HIV pathogenesis [25, 26]. Despite this potential adverse effect, early ART may limit chronic immune activation [15] and decrease the size and half-life of the latent reservoir established early in infection in infants [27].

With respect to the timing of viral rebound post ATI, there is a natural trade-off with regard to viral rebound between development of a neutralizing antibody response and the establishment of the latent reservoir, thought to be the source of viral rebound following treatment suspension. The dynamics of the latent reservoir are expected to be different between adults and children, since the infection in infants occurs in the setting of a developing T-cell repertoire and high viral replication, as well as a deficient cytotoxic T cell activity [28–33]. In infants, the latent reservoir is more homogeneous than in adults as it has not been selected by CTL and other immunologic pressures [15, 34]. Furthermore, in adults, the latent reservoir comprises mainly of memory cells. However, infants lack in memory cells and develop them in the first decade of life, reaching approximately 50% of the T-cell population [35]. Like adults, HIV-1 persists in resting CD4+ T lymphocytes in children treated with prolonged ART [36]. Despite these differences, the latent reservoir is an important factor in the eradication of pediatric HIV-1, just as it is with adult HIV.

Limiting viral antigen exposure early may be desirable, but it could affect the development of adaptive immune responses and influence disease progression if ART is suspended [37]. Understanding the differences in the virologic and immunologic aspects of pediatric HIV may inform innovative targeted strategies that may be safe for children, even if detrimental in adults [37, 38].

In this study, we are interested in disentangling the effect of antibodies and latent reservoir size in pediatric HIV after treatment interruption. We aim to understand how ART initiation time sets the stage for the trade-offs in latent reservoir establishment and neutralizing antibody development and how these in turn affect post-rebound dynamics. We use experimental data from an infant nonhuman primate Simian/Human Immunodeficiency Virus (SHIV) infection model that mimics breast milk HIV transmission in human infants. Infant rhesus macaques (RMs) were orally challenged with SHIV.C.CH505 375H dCT 4 weeks after birth and ART was initiated late at 8 weeks post-infection (wpi), intermediately at 2 wpi and early at 4-7 days post-infection (dpi). In addition to regular viral load measurements, longitudinal neutralizing antibody responses as well as the latent reservoir size were assessed after ART interruption [39–41]. We use mathematical modeling to investigate how the timing of treatment initiation affects the establishment of the latent reservoir and the development of neutralizing antibodies, and how these, in turn, affect rebound dynamics. We use the standard viral dynamics model [42, 43] fitted to viral load measurements using a nonlinear mixed effects (NLME) framework with neutralizing antibody responses and latent reservoir as covariates. Using this approach, we show that strong neutralizing antibody responses are associated with increased viral clearance and decreased viral infectivity but decreased death rate of infected cells. In addition, the absence of detectable latent reservoir is associated with more robust immune responses. Overall, we find that rebound times are decoupled from downstream dynamics, as the amount of time it takes for an RM to show sustained detectable viremia is not related to the growth rate of the virus.

## 2 Methods

In the following sections, we will describe the experiments, as well as the data we obtain. We follow with a description of the model, and the way we fit the model to the data. Finally, we provide a description of the way we compare models.

### 2.1 SHIV infection data in infant macaques

Ten infant *Rhesus macaques* (RMs) were orally challenged with SHIV.C.CH505 four weeks after birth. Antiretroviral treatment (ART) was initiated within 8 weeks post-infection (wpi) and subjects were followed for viral rebound, defined as sustained detectable viremia. Viral load measurements were taken regularly, every 2-3 days. The experiment was replicated twice with different treatment initiation times: ten animals started treatment at 2 wpi (intermediate treatment group) and another 10 RMs started 4-7 days post-infection (dpi) (early treatment group) (Fig 1). In addition to viral load measurements, the potency was neutralizing antibodies against autologous virus envelope glycoprotein gp120 was assessed by a TZM-bl assay [44] at day 0 post-ATI. The latent reservoir size was estimated with Intact Proviral DNA assay (IPDA) [45] also at day 0 post-ATI. The experiment for the late treatment group is described in detail in [40, 41].

**Figure 1:**
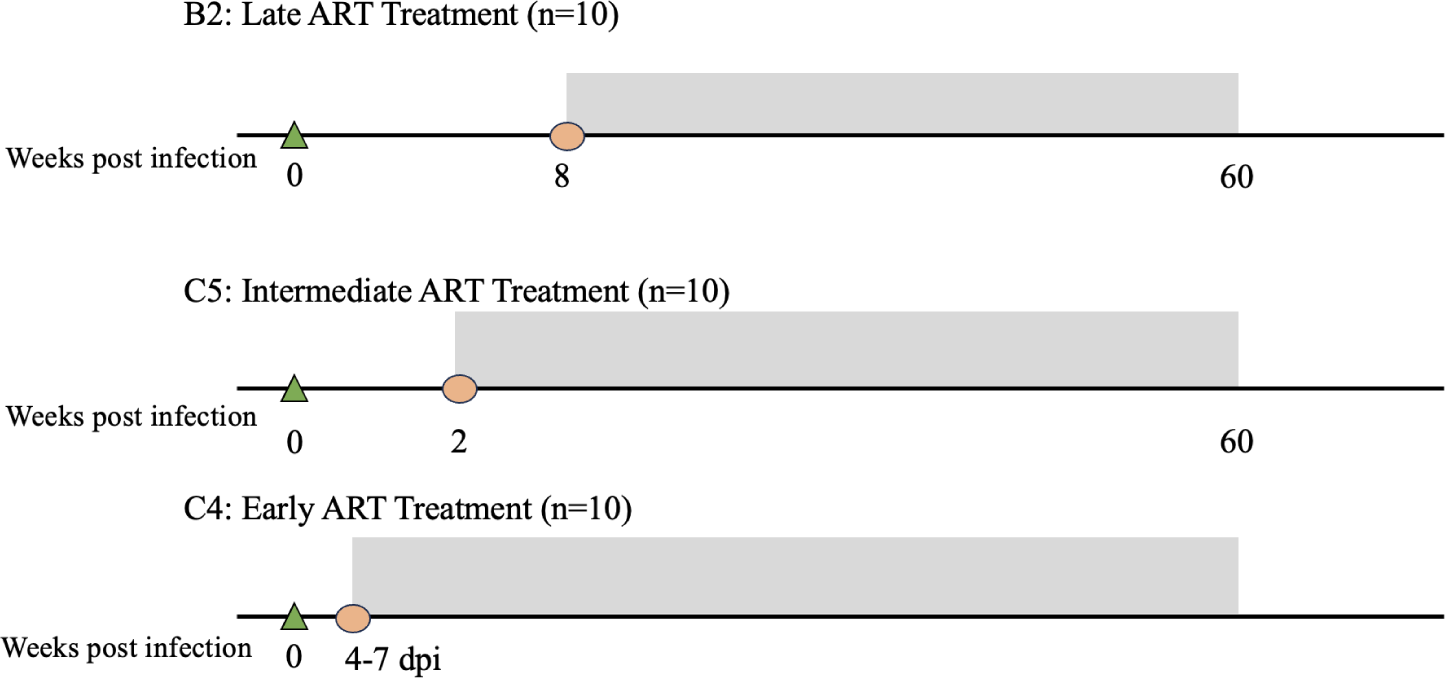
Description of the experiments. Study schematic showing ART timing (grey shaded area) and tissue collection for the three treatment groups.

We calibrate our model using the following experimental measurements: 1) viral load measurements, 2) potency of antibodies neutralizing autologous virus envelope glycoprotein gp120 measured by a TZM-bl assay [44] and 3) the latent reservoir size assessed with Intact Proviral DNA assay (IPDA) [45]. A summary of the data we use can be found in Table 1.

**Table 1:** Summary of the experimental data.

|  | **Late** | **Intermediate** | **Early** |
| --- | --- | --- | --- |
| **Viral load** | 9/10 rebounded<br>median 18 days | All rebounded<br>median 17 days | 7/10 rebounded<br>median 84 days |
| **Neutralization efficacy** | Detectable for all;<br>high for 5/10 subjects | Undetectable | Undetectable |
| **IPDA** (intact genomes/ $10^6$ CD4+ T cells) | Detectable for all subjects | Detectable for 5/10 subjects | Undetectable |

### 2.2 The model

In our study, we use the “standard” viral dynamics model, initially devised to quantify initial approximations of core within-host parameters, such as the viral clearance rate and the infected cell lifespan [42] and to investigate viral decay during treatment [43]. The model consists of three compartments– uninfected cells, infected cells, and free virus. Target cells, *T*, are produced at rate *λ*, die at rate *d* and become infected at rate *β*. Infected cells, *I*, die at a rate *δ* and produce virions at rate *p*. Free virus, *V*, is cleared from the blood at rate *c* (Fig 2). This model is succinctly expressed through the following set of differential equations:

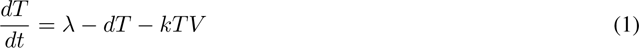

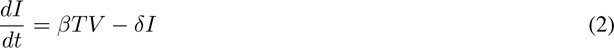

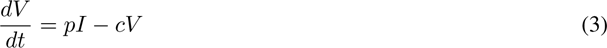

**Figure 2:**
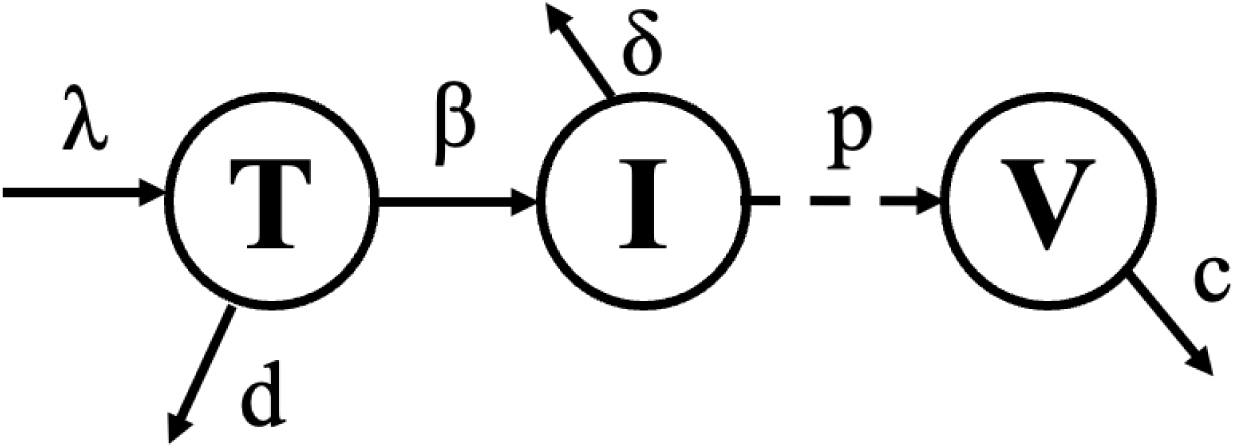
Diagram of the standard dynamics model. Diagram of the standard viral dynamics model. Target cells, *T* are sourced at rate *λ*, die at rate *d* and through contact with the virus become infected at rate *β*. Infected cells *I* die at rate *δ* and produce virions at rate *p*. Finally, free virions are cleared at rate *c*.

The standard model is a target cell-limited model, meaning that the viral load trajectory is constrained by the availability of target cells. It has been shown to capture the dynamics of early infection [42, 43, 46] (2). Our aim is to capture broad differences among groups. For that reason, we focus on a simple and parsimonious model– the standard model fulfills both criteria [47]

We rescale *T* and *I* by *T*_0_, where *T*_0_ is the initial number of target cells per mL. It should be noted that viral production rate *p* and *T*_0_ structurally unidentifiable and therefore will be calculated as a composite parameter. We also assume that prior to infection target cells are at equilibrium and therefore, we set *λ* = *d · T*_0_ [46]. The only parameter we fix *d* = 0.01 per day [46]. We estimate the remaining four parameters: *β*, *δ*, *p*, *c* of the model, as well *t_start_*, which is the onset of exponential viral growth, for total of five parameters. We assume that starting at treatment interruption, there is a stochastic phase during which viral loads are below the detection threshold. Once that threshold is crossed, the virus grows exponentially and dynamics become deterministic, thus captured by the standard model. *t_start_* captures the time frame between treatment interruption and the start of exponential viral growth.

### 2.3 Motivating Calculations

We first fit the standard viral dynamics model to each treatment group separately, to investigate if there are differences in the dynamics between the three treatment groups. We fit the standard viral dynamics to viral load measurements for all three treatment groups, in a nonlinear mixed effects (NLME) framework. We are particularly interested in elucidating the differences between groups. NLME allows us to capture both the differences between pre-defined groups as well as the variability among study participants.

We find that the main difference between the treatment groups lies in the viral clearance rate: for the late treatment group, the population value of the viral clearance rate is double compared to that for the intermediate and early groups (Supplementary Table 1). In addition, mass action infectivity is approximately one order of magnitude lower for the early treatment group. Finally, the start of exponential viral growth is similar between the late and intermediate treatment group and much higher for the early group (Supplementary Table 1), which is also consistent with rebound times (Table 1).

### 2.4 Model Fitting

In the previous section we compared the three treatment groups and found that they differ in their viral dynamics. Now, we seek to investigate if the differences observed among treatment groups can be explained by the latent reservoir or neutralizing antibodies instead. We incorporate the effect of the latent reservoir and neutralizing antibody responses by allowing parameter values of the standard model to differ in the presence or absence of detectable latent reservoir and in the presence or absence of potent neutralizing responses. Any of our estimated parameters - viral infectivity *β*, viral clearance *c*, infected cell death *δ*, scaled viral production *pT*_0_, or onset time exponential viral growth *t_start_* - may vary with latent reservoir size, neutralization efficacy, both, or neither. We test models with all possible combinations of our five model parameters having different values either in the presence of detectable latent reservoir or potent neutralizing antibodies or both simultaneously (Fig 3. Therefore, we will identify which standard model parameters are predicted to differ in the presence of the two biomarkers and generate hypotheses to describe different infection mechanisms associated with neutralizing antibody responses and latent reservoir size.

**Figure 3:**
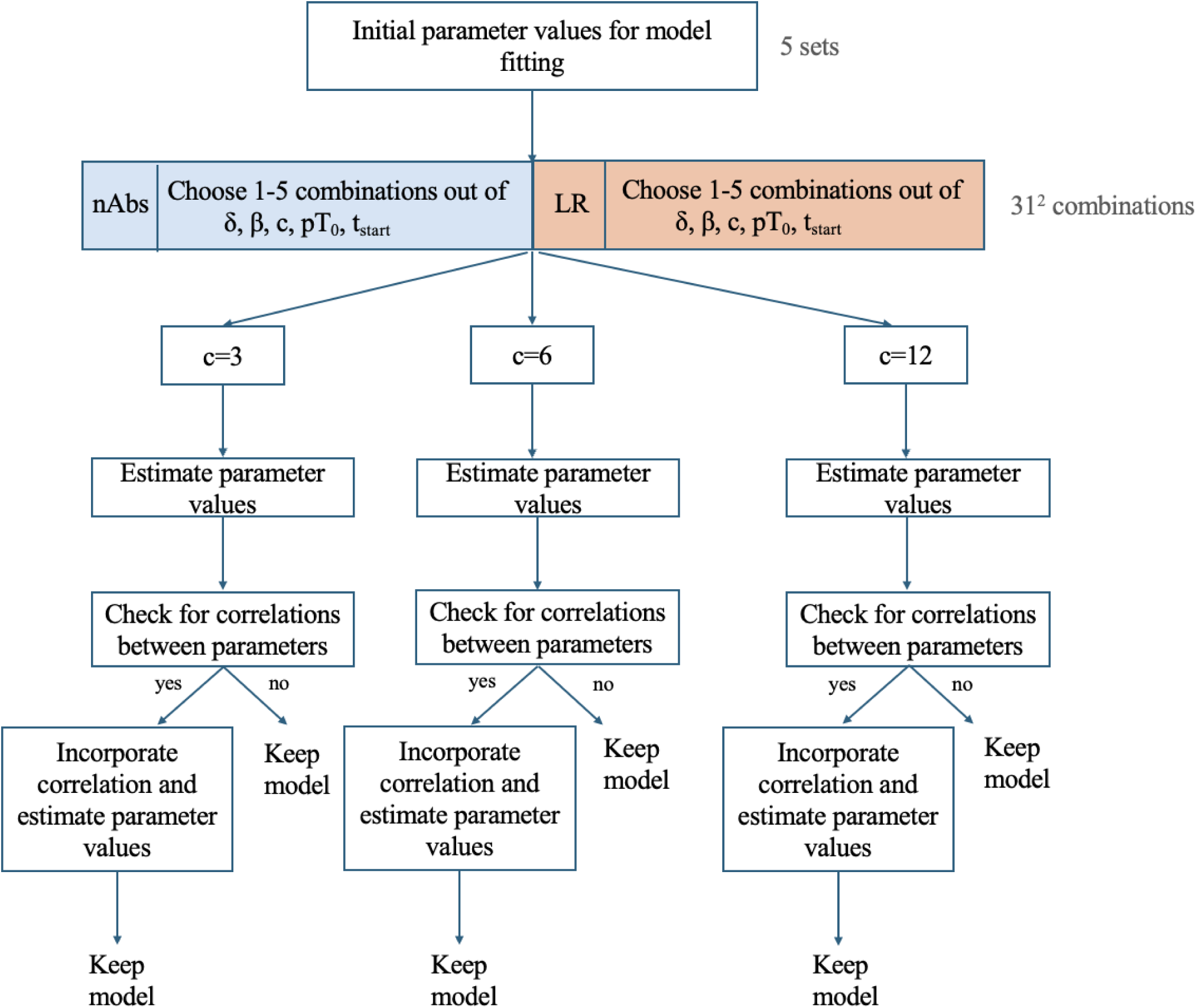
Model Fitting Schematic. Schematic of the approach we follow for model fitting and types of models tested.

We fix the median viral clearance rate to *c* = 3, 6, 12 and 24 day*^−^*^1^ [43, 46, 48], while still allowing for inter-individual variability. To better explore the parameter space, we test each model using the best initial parameter value guess along with four sets of initial parameter values selected randomly from an interval around the original value. We add correlations between random effects in pairs of parameters when ANOVA tests indicate significance at the 0.01 significance level. We run all models in *Monolix* through the *lixoftConnectors* package in *R*. We test a total of 25416 models.

### 2.5 Model Selection

Given the abundance of models, we do not seek to select a single model that best fits our data. Instead, we focus on an ensemble of models that are comparably successful in explaining the observed data. We determine these models using the following criteria, developed in Mainou et al. [49]:

1. Akaike Information Criterion (AIC): we consider models whose AIC is within 10 points from the lowest AIC (Fig 4).
2. Quantitative Predictive Check (QPC): QPC is a reproducible quantification of VPC (Supplementary Information 1.4).
3. Stability of parameter estimates: we evaluate the % relative standard error (RSE), which is a statistical measure that quantifies the increase in the standard error of a parameter estimate. To assess the quality of a model RSE, we devise a metric titled *S* defined as the sum of RSE values above 50% divided by the number of parameters estimated.

**Figure 4:**
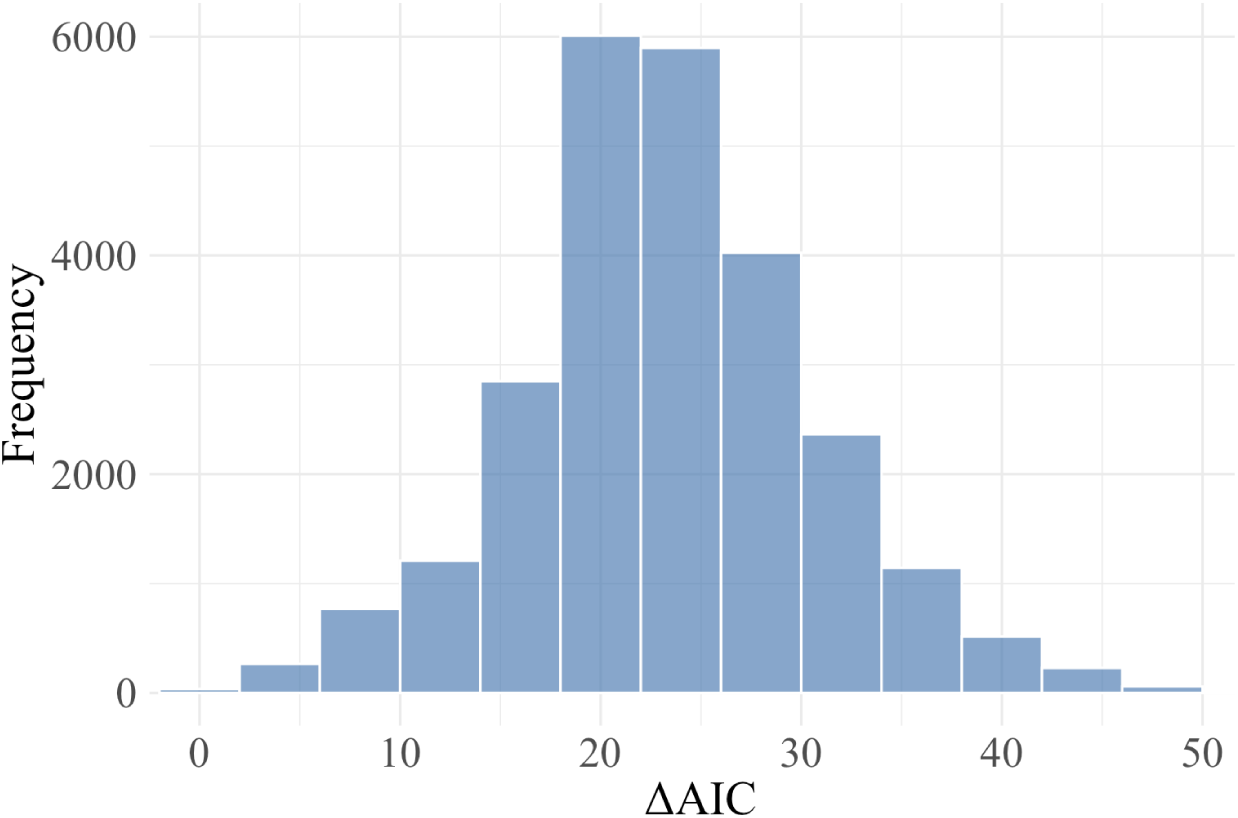
Distribution of Δ*AIC*. Histogram of AIC values relative to the lowest AIC of all the models tested. We reject models with Δ*AIC >* 10.

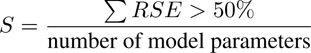

We select models under different sets of evaluation criteria that vary the stringency of one of the three evaluation metrics:

a. Δ*AIC <* 8, *QPC <* 15 and *S <* 250.
b. Δ*AIC <* 10, *QPC <* 15 and *S <* 250.
c. Δ*AIC <* 10, *QPC <* 14.5, and *S <* 250.

*S* = 250 corresponds to 10% of the highest *S* value, whereas *QPC* = 15 and *QPC* = 14.5 correspond to the 10*^th^* and 5*^th^* percentile respectively. We test the sensitivity of this combination of metrics and we acknowledge that selected models could differ if we impose different restrictions on *S* and *QPC* values (Supplementary Figs 3 and 4).

We find a set of nine models is consistently selected across all three types of model selection we employ. These models have certain common features: strong neutralizing antibody responses are a categorical covariate for the viral clearance rate (*c*), the death rate of infected cells (*δ*) and the start of exponential viral growth (*t_start_*), but not for mass-action infectivity (*β*). Similarly, latent reservoir is a covariate predominantly for *t_start_*, followed by *p · T*_0_ by *c* and *δ*. These observations are qualitatively consistent across all three types of model selection (Supplementary Fig 2A).

## 3 Results

We fit the standard model incorporating strong neutralizing antibody responses and detectable latent reservoir by allowing the population-level parameter values to differ in the presence/absence of these two biomarkers. We do not separate RMs based on their treatment group; instead we consider animals from all treatment groups together and animals are differentiated by the presence/absence of neutralizing antibodies and detectable latent reservoir. For the reader’s convenience we report parameter estimates and standard error (Table 2, as well as model fits, visual predictive check and distribution of individual parameter estimates (Fig 5) for one of the models selected. Additionally, we report below the results of a set of 15 models selected based on Δ*AIC <* 8, *QPC <* 15 and *S <* 250 (Fig 6), but the observations are consistent across types of model selection (Supplementary Fig 3).

**Figure 5:**
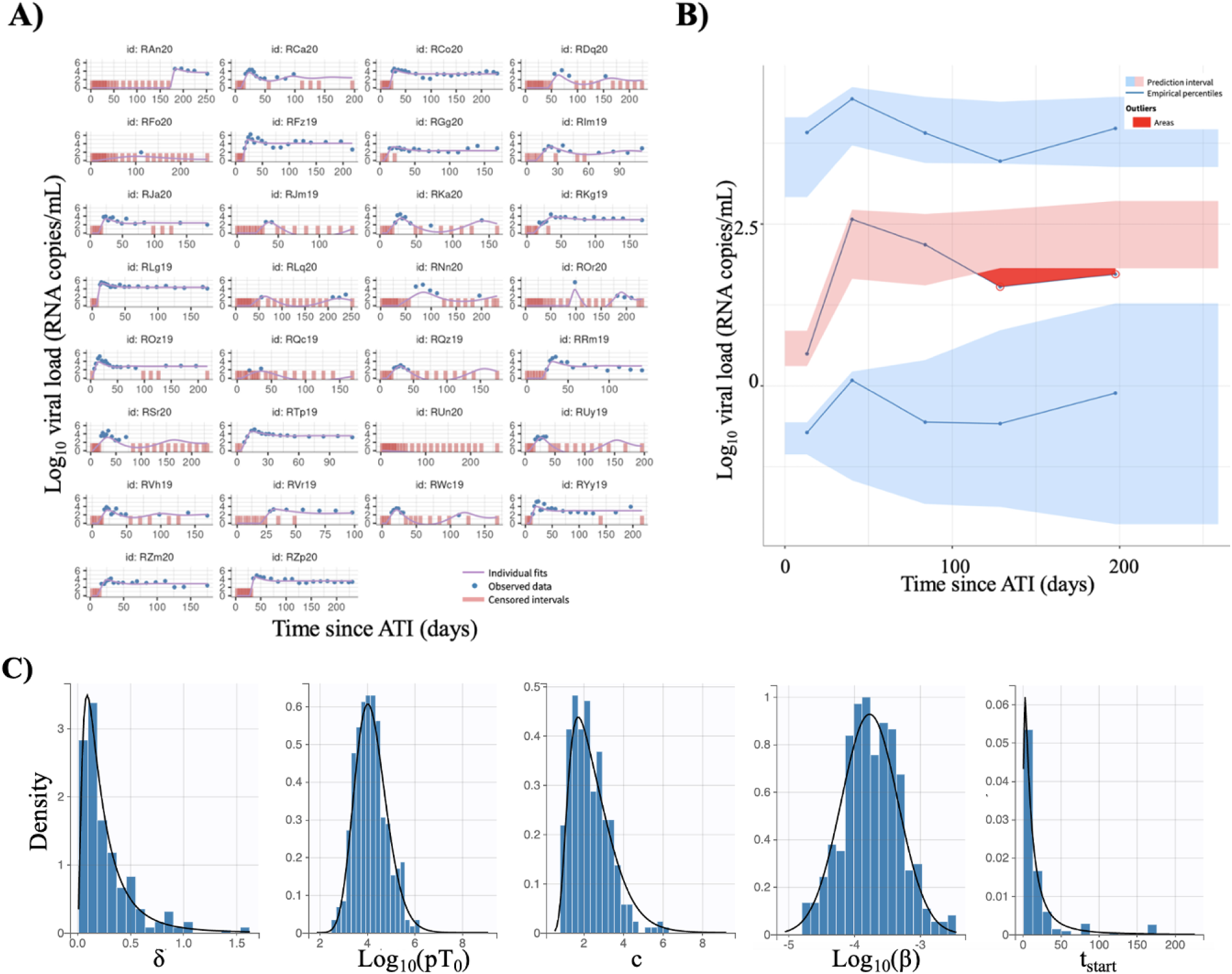
Summary characteristics of one of the models selected. Fitted curves to viral load measurements (A), visual predictive check (B) and distributions of individual parameter values (C) for one of the models selected. This particular model incorporates strong neutralizing antibody responses as a categorical covariate on the onset of exponential viral growth, detectable latent reservoir as a categorical covariate on viral clearance rate and a correlation between mass action infectivity and viral production rate.

**Figure 6:**
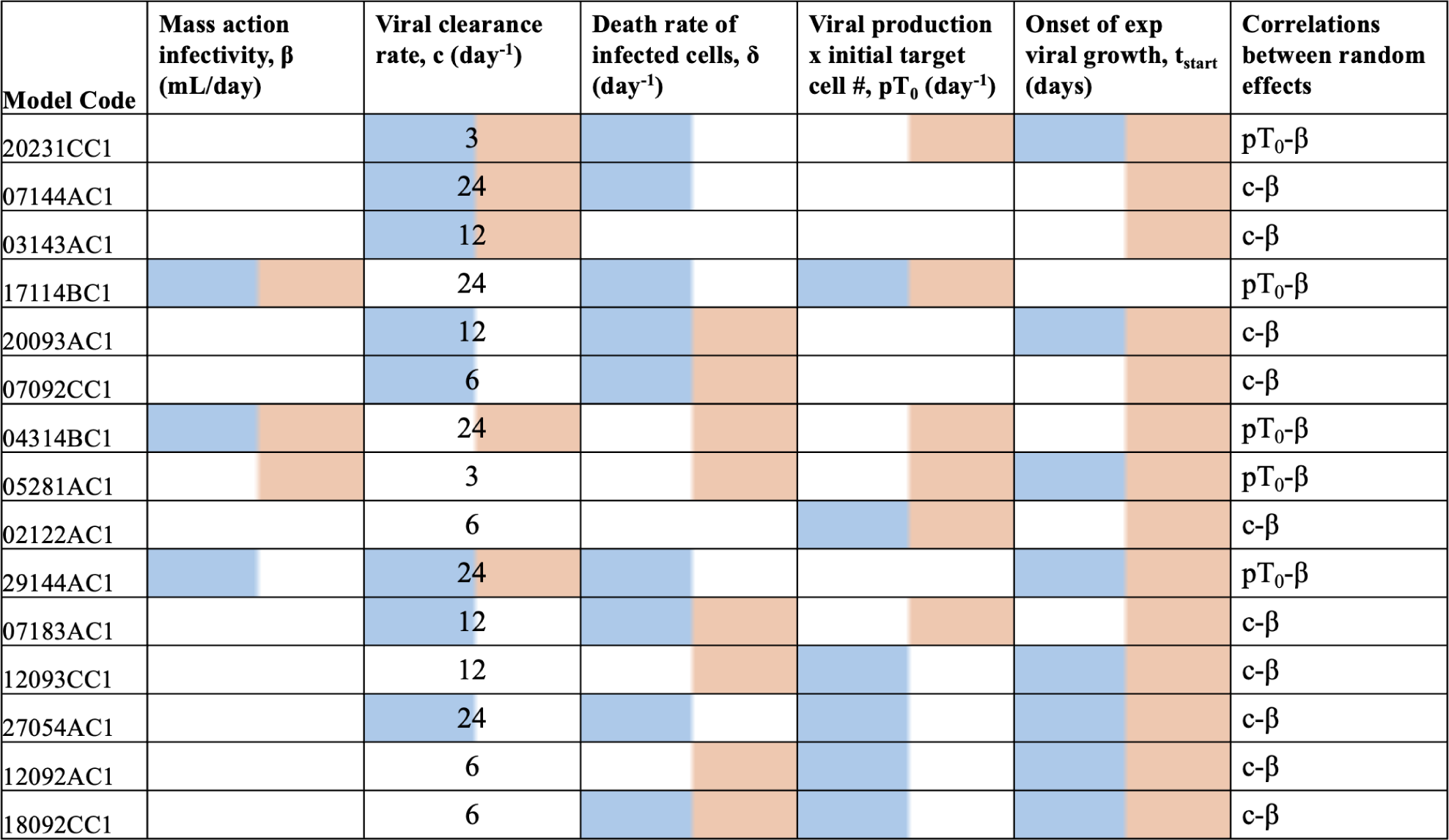
Summary of selected models. For each of the selected models, we indicate for which parameters the model assumes a difference in the median value of that parameter in the presence of potent neutralizing antibody responses (blue) or in the presence of detectable latent reservoir (orange), along with the fixed median value of the viral clearance rate *c*, and the parameters for which we incorporate correlations between random effects.

**Table 2:** Parameter values: estimates and relative in standard error (RSE) for one of the models selected (also Fig 5). This model incorporates strong neutralizing antibody responses as a categorical covariate on the onset of exponential viral growth, detectable latent reservoir as a categorical covariate on viral clearance rate and a correlation between mass action infectivity and viral production rate. the median value for viral clearance rate is fixed to 3 d*ay^−^*^1^. RSE values between 50-100% are shown in olive green and values between 100-200% are shown in orange.

|  | Value | RSE % |
| --- | --- | --- |
| Fixed effects |  |  |
| $\delta$ | 0.23 | 24.4 |
| $\log_{10}(pT_0)$ | 4.4 | 5.27 |
| $c$ | 3 | |
| $\beta_c(LR = yes)$ | -0.19 | 11.2 |
| $\log_{10}(\beta)$ | -3.92 | -100.7 |
| $t_{start}$ | 10.87 | -77.67 |
| $\beta_{t_{start}}(nAbs = yes)$ | -0.16 | 6.01 |
| Standard deviation of the random effects |  |  |
| $\delta$ | 1.02 | -126.3 |
| $\log_{10}(pT_0)$ | 0.15 | 4.53 |
| $c$ | 0.22 | 149.21 |
| $\log_{10}(\beta)$ | 0.53 | 34.78 |
| $t_{start}$ | 1.1 | 24.69 |
| Correlations |  |  |
| $\log_{10}(pT_0) - \log_{10}(\beta)$ | -0.945 | 46.45 |
| Error model parameters |  |  |
| $a$ | 0.71 | -3.43 |

### Antibodies and latent reservoir cannot explain the dynamics on their own

Models containing the presence of strong antibody responses only or the presence of detectable latent reservoir size consistently underperform compared to models including both effects by AIC (Δ*AIC >* 10.3). This suggests that either biomarker is not sufficient to explain dynamics independently. Instead, all models selected incorporate both antibody responses and the latent reservoir effects.

### Neutralizing antibodies are associated with increased viral clearance rate but decreased killing of infected cells

Below we report how many of the selected models predict a change in each of the model’s parameters in the presence of neutralizing antibodies, as well as whether this change is an increase or decrease. First, we find that the presence of strong neutralizing antibody responses is associated with a decreased killing rate of infected cells, *δ* (Fig 7A). This decrease in the average value of *δ* in the presence of neutralizing antibodies is high, approximately 55% compared to when potent neutralizing antibodies are not developed. Strong neutralizing antibodies are also associated with an increase in the viral clearance rate *c* (Fig 7A), which is expected considering the function of neutralizing antibodies [50]. This increase is very high with an average of 154%. The effect of neutralizing antibodies on mass-action infectivity is harder to parse out: for only 3 out 15 selected models, neutralizing responses are associated with a change in mass action infectivity. In two instances, neutralizing antibodies are correlated with an increased median value for mass-action infectivity, but in one model with a decreased value (Fig 7A). Viral production rate and the number of CD4+ T cells at the onset of infection (*p · T*_0_) appear as a composite parameter and therefore are estimated as one. The presence of strong neutralizing antibody responses is mostly associated with a decrease of approximately 20% in *p · T*_0_ (Fig 7A). Finally, in two out of the 15 selected models a reduction in the population-level value of *t_start_*is observed in the presence of strong neutralizing antibody responses, whereas in 7 out of 15 an increase is observed. However, the changes in *t_start_* associated with nAbs are moderate: for the 7 models the increase is on average 22%, whereas for the two models the decrease averages 10%.

**Figure 7:**
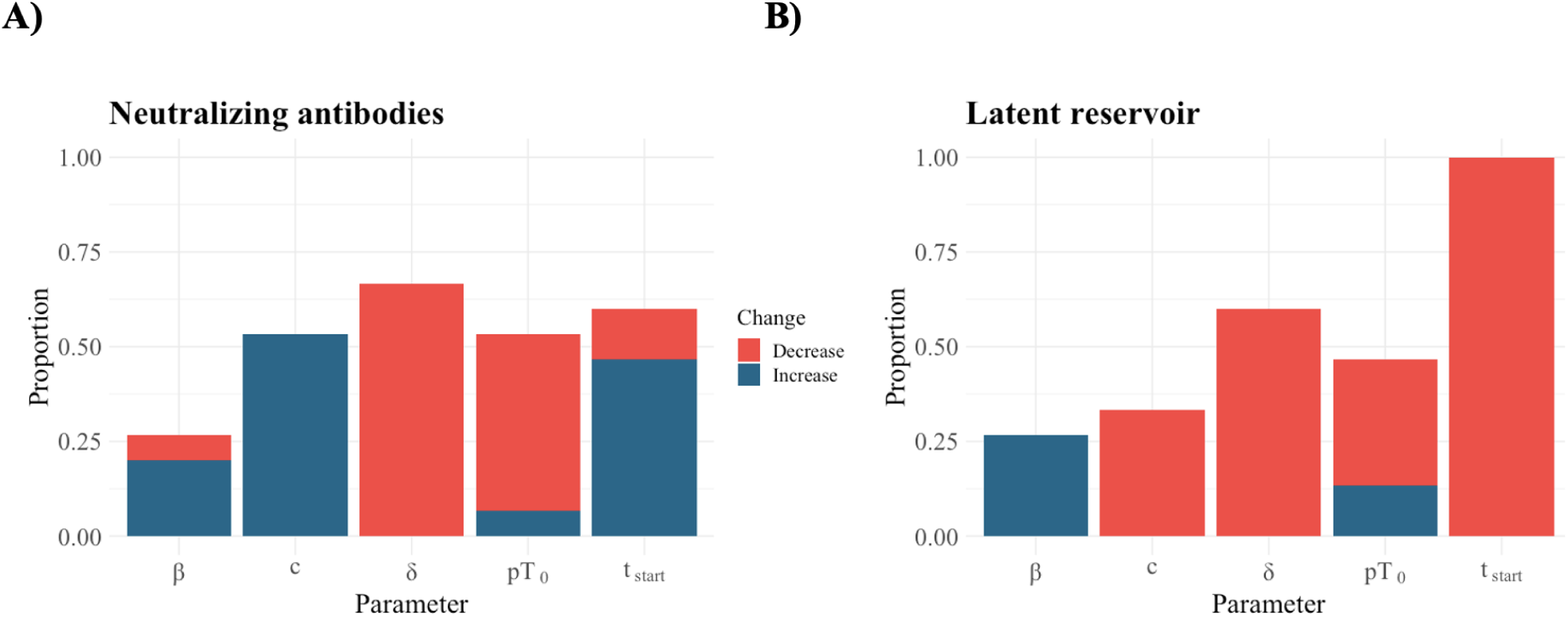
Effect of neutralizing antibodies and latent reservoir. Barplot depicting the proportion of models the population-level value of specific parameter is predicted to be different in the presence of strong neutralizing antibody responses (A) or in the presence of detectable latent reservoir (B). Blue indicates that the associated change is an increase, whereas red signifies a decrease.

### Latent reservoir is associated with less robust immune responses

Similarly, we report how many of the selected models predict a change in population-level value in each of the models parameters in the presence of detectable LR, as well as whether this change is an increase or decrease. The presence of detectable LR is consistently associated with a shorter *t_start_*, which designates the viral rebound time (Fig 7B). The effects of the LR are considerable, since the presence of detectable LR is associated with a decrease in rebound times by 72%. The presence of detectable LR is also associated with a decrease by approximately 20% in the population-level value scaled viral production rate (*pT*_0_) in approximately five of the selected models. (Fig 7B, Supplementary Fig 3B). For two models we find an increase *pT*_0_, however that increase is only slight (approximately 1.3%). Again, since the viral production rate and the initial number of target cells are non-identifiable, it is difficult to attribute the change in their values. Finally, the presence of detectable LR is associated with decreased viral clearance rate *c* (ranging between 13 *−* 74%), decreased death rate of infected cells *δ* (approximately 37%) and increased mass-action infectivity *β* (approximately one order of magnitude) (Fig 7B). A recurring pattern emerges: detectable LR is linked to an underdeveloped immune response. We posit that this underdeveloped immune response permits larger LR sizes. The intermediate treatment group shows no antibody responses but detectable LR, whereas RMs treated the earliest, show no antibody responses and no detectable LR. This lack of detectable LR is attributed to the timing of treatment initiation, immediately upon seroconversion, which vastly limits the establishment of the LR.

Overall, we find that the associations of the LR size and parameter changes among are more consistent among selected models compared to the effects of potent neutralizing antibodies.

### The model predicts that viral rebound times are decoupled from downstream dynamics

We observe that the time of onset of exponential viral growth is statistically higher for RMs that did not develop potent neutralizing antibodies nor detectable latent reservoir compared to the RMs that developed either (Supplementary Table 8 and Supplementary Fig 5). We estimate the viral growth rate (Supplementary Text) for each RM in all selected models. We find no correlation between the time of onset of exponential viral growth (*t_start_*) and the viral growth rate (*r*) (Pearson and Spearman correlations p-values*>* 0.01, Supplementary Table 8). In addition, we find no linear trend between *t_start_* and all model parameters (Supplementary Table 10 and Supplementary Fig 6).

## 4 Discussion

This study used a viral dynamics model with a nonlinear mixed-effects framework to examine how treatment initiation time affects the latent reservoir and neutralizing antibody responses and how these, in turn, post-rebound dynamics in SHIV-infected infant RMs. Unlike most studies, we focused on post-rebound dynamics rather than just rebound times. The model is able to accurately depict rebound dynamics, but underestimates peak viremia [47]. We found that strong neutralizing antibody responses led to increased viral clearance but reduced death rates of infected cells, while the absence of detectable latent reservoirs was linked to a stronger immune response. Our analysis also shows that the timing of the onset of exponential viral growth is not related to the viral growth rate or any other parameters in our model, though our sample size was small. This implies that the dynamics between treatment interruption and the start of viral rebound are independent of what happens after the rebound begins, suggesting that strategies to delay rebound could differ from those needed to manage post-rebound growth. Viral evolution might explain these findings, as studies on broadly neutralizing antibodies (bnAbs) indicate that rebounding viruses can develop resistance to bnAbs used as monotherapies [51].

Importantly, our results highlight that models incorporating both neutralizing antibody responses and latent reservoir data provide a more comprehensive understanding of post-rebound dynamics, suggesting that a tailored approach might be necessary for medical interventions. Although latent reservoirs showed a consistent impact across models, indicating a clear role in post-rebound dynamics, the effect of neutralizing antibodies was less straightforward, suggesting a dependency on other factors. This variability underscores the complexity of immune responses and indicates that a broader range of datasets and more intricate models may be required to fully grasp these dynamics.

Our results highlight the role of both antibody responses and the latent reservoir in influencing rebound dynamics. We find that stronger neutralizing antibody (nAb) responses correlate with a reduced rate of infected cell death. This could indicate a disruption in the function of CD4+ T cells, which are crucial for activating cytotoxic T cells (CD8+ T cells), the primary cells responsible for eliminating infected cells. A reduction in CD4+ T cell activity may lead to decreased CD8+ T cell-mediated killing, ultimately affecting rebound dynamics [52]. Further studies are needed to better understand these mechanisms. We also observe that neutralizing antibodies are associated with an increase in mass action infectivity for a very small subset of selected model, even though the increase is slight. This contrasts with previous mathematical modeling work using human adult data that demonstrated a correlation between anti-gp41 antibodies and a decay in viral infectivity [53]. In this study we use anti-gp120 antibodies and we hypothesize that this increase points to immune exaustion. Similarly, the increase in rebound times in the presence of neutralizing antibodies could be attributed to neutralization escape, as it would be expected that the presence of neutralizing antibodies is associated with a prolonged rebound time.

Finally, we observe a decrease in the product of viral production rate and initial number of target cells in the presence of potent antibody responses. Since *p* and *T*_0_ are not identifiable, we cannot determine which parameter accounts for this decrease. However, we hypothesize that this decrease can be attributed to either parameter. Regarding *T*_0_, in pediatric HIV, CD4+ T cells do not recover to pre-infection levels under ART [24, 54, 55]. Viral production rate could also be lower at later stages of the infection due to non-cytolytic effects of CD8+ T cells with recent modeling works providing support for a dual function of CD8+ T cells in HIV-1 infection [56].

Overall our results indicate that latent reservoir is associated with stronger immune responses and increased control. First we observe that all models predict a decrease in the onset of exponential growth *t_start_* in the presence of detectable latent reservoir. The start of exponential viral growth designates viral rebound time. Our finding is consistent with the central hypothesis that the activation of latently infected cells drives viral rebound [57–60], as well as with previous modeling results of the same dataset [61]. The latent reservoir size is also associated with changes in immune-related parameters consistent with stronger immune responses, both cellular and humoral. Indeed, this has been observed in a set of adult post-treatment controllers for whom the total proviral genome is correlated with NK cell and T cell activation [62]. We also observe that the presence of latent reservoir is mostly associated with lower average value of the product of the viral production rate and the initial number of target cells *p · T*_0_. Again, since these two parameters are non-identifiable, it is difficult to attribute the change in their values. We do hypothesize, though, that the decrease in *p·T*_0_ could stem from a reduction in CD4+ T cells due to the infection-induced damage of CD4+ T cell responses [24, 54, 55]. In the two models for which an increase in the product is observed, *T*_0_ could decrease, but the viral production rate could also increase, making up for a reduced *T*_0_. We hypothesize that the positive association between the presence of detectable latent reservoir and increased viral production rate could be attributed to a more intense infection; producing more virions per unit time could lead to the infection of more cells, some of which contribute to the latent reservoir.

An important limitation of our study is that the effects of the latent reservoir and neutralizing antibodies are associations with the parameters of interest. Even though this provides a simple framework to explore patterns in the relationship between our biomarkers and key viral kinetics parameters, sometimes it is difficult to disentangles those relationship. For example, we find in the 8 models that predict an effect of strong neutralizing responses 6 of them predict an increase in the viral clearance rate, whereas 2 predict a decrease. It may be a small number of selected models that predict a decrease; however this has important implications in determining the possibility of the development of neutralization resistance or immune escape. Perhaps a more explicit way to model the relationship between antibodies and viral clearance rate, such as a Hill function of antibody responses multiplied by the viral clearance rate, could shed better light on the relationship between the two. Another limitation of our model is the consistent misspecification of the nadir and subsequent viral increase (Fig 5B). That region of the viral load trajectory shows great variability with RMs showing both detectable and undetectable measurements. To capture that region, we used the standard model incorporating density-dependence in the death of infected cells, which has been shown to capture viral decay better [47, 48, 63]. However, this model was not able to improve the goodness of fit, suggesting that there are mechanisms not included and needed to explain this variability.

Overall, our findings indicate that for pediatric HIV infections, early treatment initiation is crucial for achieving the best outcomes, especially when treatment interruptions occur. The influence of the latent reservoir on rebound dynamics is clear and predictable. In contrast, immune responses are complex and their effects are more difficult to discern. As such, delaying treatment to encourage the development of adaptive immune responses is not advisable, as the impact of these responses is less certain compared to the well-documented effects of a larger latent reservoir. Instead, it is advisable to start treatment early to minimize the size of the latent reservoir, with the potential to later augment immunity through emerging immune-based therapies (reviewed in [15]). This recommendation is supported by case studies in both adults and infants. For example, the “Mississippi baby” was treated aggressively only 30 hours after birth showed no detectable viremia for up 28 months after treatment interruption [17, 18]. Recent studies in children also support that very early treatment may enable sustained remission [64]. In adults, a pooled analysis of 6 AIDS Clinical Trials Group ATI studies revealed that participants who started ART during acute/early HIV infection has significantly delayed rebound [16]. However, this is not fully supported by other experimental findings in adult RMs. An analysis of combined data from 10 independent studies on adult SIV-infected RMs found that up to 3 weeks post-infection, each day of delay in treatment initiation is associated with lower peak viremia and setpoint. After 3 weeks post-infection delaying treatment is associated with higher peak and setpoint [65].

Despite limitations and simplifications, this study examines post-rebound dynamics by considering infant RM-specific information on latent reservoir size and neutralizing antibody potency in postnatal SHIV infection. Our results offer insight into the post-rebound dynamics in pediatric HIV infections and underscore the critical importance of early treatment initiation in pediatric HIV infections, shedding light on the pivotal role played by antibody responses and latent reservoir size in shaping post-rebound dynamics.

## Acknowledgements

We would like to thank Dr. Ephraim Hanks for the helpful discussions on developing the Quantitative Predictive Check. J.M.C. acknowledges the support of the National Science Foundation (grant no. DMS-1714654) and National Institutes of Health (grant nos. R21-AI143443-01A1 and R01-OD011095). E.M. acknowledges the support of the National Science Foundation (grant no. DMS-1714654). G.M.S acknowledges the support of the National Institutes of Health (grant no. R01 AI160607). C.C. acknowledges the support of the National Institutes of Health (grant no. R25AI140495). S.R.P. acknowledges the support of the National Institutes of Health (grant no. P01-AI131276-05).

## Data and computational code availability

The data utilized in this study originates from experimental research. For access to these data, please contact the Principal Investigator of the study, Dr. Ann Chahroudi, directly. All analysis code, along with a simulated test dataset are available at https://github.com/elliemainou/reboundPO1-nlme.

## Notes

### Competing Interest Statement

Dr Conway has served as a consultant for Excision BioTherapeutics and Merck. Dr. Permar serves a consultant for Moderna, Merck, Pfizer, GSK, Dynavax, and Hoopika on their CMV vaccine program and has led a sponsored program with Moderna and Merck on CMV vaccines.

