## Supplementary material for "Assessing the impact of autologous neutralizing antibodies on rebound dynamics in postnatally SHIV-infected ART-treated infant rhesus macaques": Rebound P01 Supplementary Figures.docx

Supplementary Information

**Section 1: Supplementary Text**

1. Derivation of rescaled standard viral dynamics model

The standard viral dynamics model consists of the following set of differential equations:

$$\frac{dT}{dt}=\lambda-dT-\beta TV$$

$$\frac{dI}{dt}=\beta TV-\delta I$$

$$\frac{dV}{dt}=pI-cV$$

We set that $\lambda=dT_{0}$ [3] and assume that $Ť=\frac{T}{T_{0}}$ $\to$ $T=ŤT_{0}$ and $Ĩ=\frac{I}{T_{0}}\to$ $I=ĨT_{0}$. Substituting to the original equations,

$$\frac{dŤT_{0}}{dt}=dT_{0}-dŤT_{0}-\betaŤT_{0}V$$

$$\frac{dĨT_{0}}{dt}=\betaŤT_{0}V-\deltaĨT_{0}$$

$$\frac{dV}{dt}=pĨT_{0}-cV$$

Simplifying the system and renaming $Ť$as T and $Ĩ$ as I, we obtain the final system:

$$\frac{dT}{dt}=\lambda-dT-\beta TV$$

$$\frac{dI}{dt}=\beta TV-\delta I$$

$$\frac{dV}{dt}=pT_{0}I-cV$$

Since the p and T_0_ appear as a product only, these parameters are structural unidentifiable and thus estimated as one.

2. Standard viral dynamics model incorporating density-dependent decay of infected cells

Fitting the standard viral dynamics [1-3] model to our viral load measurements, we observed a consistent misspecification around the nadir in the visual predictive check. To remedy that, we tried instead a version of the standard model that incorporates density dependence in the decay of infected cells. We report parameter estimates, the rise in standard error (Supplementary Table 2), as well as the visual predictive check and a sample of model fits (Supplementary Fig 2). We did not perform an exhaustive search for the best version model, but we report the results for one of the best runs.

3. Relative Standard Error (RSE)

The % RSE is a statistical measure that quantifies the increase in the standard error of a parameter estimate when comparing two different models or conditions. The RSE is a useful metric that helps assess whether changes in the model's complexity or data have a substantial impact on the reliability of parameter estimates. RSE values greater than 50% warrant caution regarding the stability of parameter estimates. We exclude models where standard error for at least one estimated parameter could not be calculated. To assess the quality of a model RSE, we devise a metric titled S defined as the sum of RSE values above 50% divided by the number of parameters estimated.

$$S= \frac{sum RSE>50\%}{number of model parameters}$$

For example, for the model estimates in Table \ref{tab:parametervalues}, the estimated $S$ value would be

$$S= \frac{100.7+77.67+126.3+149.21}{13}=34.91$$

4. Quantitative Predictive Check (QPC)

The Visual Predictive Check (VPC) is a graphical method used in statistical modeling and outputted by Monolix to assess the goodness-of-fit and predictive performance of a model (Supplementary Fig 1). It's a valuable tool for evaluating how well a model captures the observed data and how well it can predict future observations. However, since it is a graphical representation, it can be difficult to assess the differences between VPC figures of different model. For this reason, we develop the QPC, to quantify the VPC.

To assess the predictive ability of each model, we assess the ability of a model to capture the viral load trajectory of each study participant.

We have the general model:

$y_{ij}=f\left( t_{ij}, \psi_{i} \right)+\alpha\varepsilon$

Where $y_{ij}$ is the j^th^ observation of the i^th^ individual, *f* is the structural model (i.e. standard viral dynamics model) based on the time point $t_{ij},$ and the parameter values for that i^th^ individual, $\psi_{i}$. $\alpha$ is the constant residual error model parameter and $\varepsilon$ is the residual error, normally distributed with mean 0 and standard deviation 1.

For each study participant we simulate the model 500 times using that individual’s parameter values and adding the residual error $\varepsilon$ (grey shaded area in Supplementary Figure 2). From those 500 simulations, we estimate the 10^th^ and 90^th^ percentile for each time point. Then, we calculate the sum of the distances of all viral load measurements that fall outside of the 80^th^ prediction interval to the prediction interval ($\sum|viral load measurement-closest viral load prediction |$). We evaluated additional metrics, but this metric allows us to also assess the predicted deviation of a measurement to the prediction interval. The metrics we evaluated are the following:

a. Sum across RMs the proportion of viral load measurements that fall outside the prediction interval.
b. Count the number of RMs for whom the proportion of viral load measurements that falls outside the prediction interval is above a specified threshold. This metric was not sensitive enough, as it allowed for multiple models to have equal QPC values, even though the VPC varied.
c. Count the total number of points, across all RMs, that fall outside the prediction intervals.
d. For all points that fall outside of the prediction interval, sum their distance to the prediction interval.
e. Take the square root of the sum of all squared distances for measurements that fall outside the prediction interval.

$$\sqrt{\sum{(viral load measurement-closest viral load prediction)}^{2}}$$

Of all five metrics we evaluated, we use the sum of distances (d) because we find that a metric that also accounts for the deviation from the prediction interval is more appropriate. The resulting metric ranges between 13.82 and 18.42. It should be noted though that the viral curve around the nadir is consistently misspecified. To remedy this, we used the standard viral dynamics model incorporating a density-dependent decay of infected cells, which has been shown to better capture post-peak viral decay [4, 5], but model results were quite poor (Supplementary Table 2 and Supplementary Fig 1).

5. Estimation of viral growth rate

The viral growth rate is the dominant eigenvalue of the system. At the beginning of the infection, we assume that at the beginning of the infection $T=T_{0}$. Therefore,

$\frac{dI}{dt}=\beta TV-\delta I$= $\beta T_{0}V-\delta I$ and

$\frac{dV}{dt}=pI-cV$*.*

This can be rewritten as ( $\begin{matrix} I \\ V \end{matrix}$ )'= $\left( \begin{matrix} -\delta& {\beta T}_{0} \\ p & -c \end{matrix} \right)$ ( $\begin{matrix} I \\ V \end{matrix}$ ), where A= $\left( \begin{matrix} -\delta& {\beta T}_{0} \\ p & -c \end{matrix} \right)$

Using the determinant,

$\det\left( A-rI \right)=0$ $\to$

$$\left( -\delta-r \right)\left( -c-r \right)-\beta pT_{0}=0\to$$

$$r^{2}+\left( c+\delta\right)r+c\delta-\beta pT_{0}=0$$

Therefore, the viral growth rate is $r=\frac{-\left( c+\delta\right)+\sqrt{(c-\delta)^{2}-4\beta pT_{0}}}{2}$

**Section 2: Supplementary Figures**

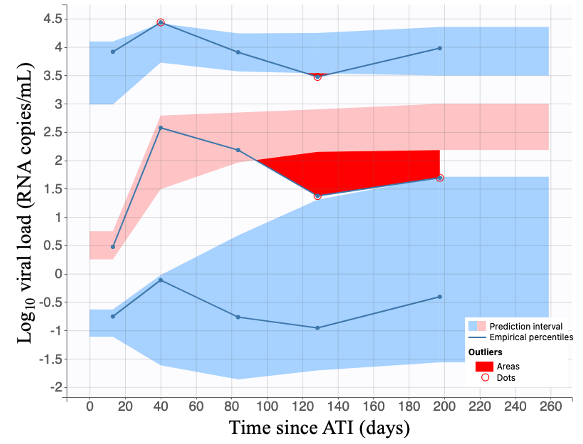

**Supplementary Fig 1**: Visual predictive check derived for the standard model that incorporates density-dependence in the death of infected cells. The solid lines are the empirical percentiles (10^th^, 50^th^ and 90^th^) and the shaded areas are the prediction intervals. Areas where the empirical percentile falls outside of the respective prediction interval are signified in red. Overall, the areas of misspecification are worse compared to multiple of the standard models selected in our analysis.

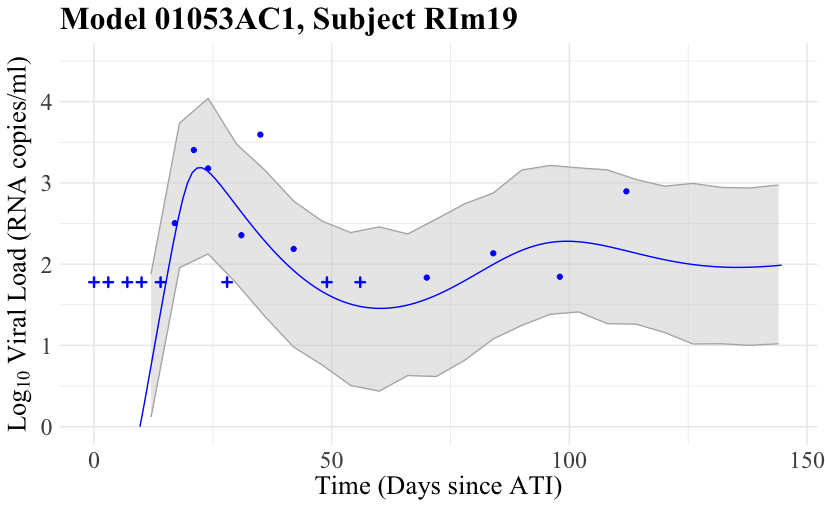

**Supplementary Fig 2**: Example of a Quantitative Predictive Check for model 01053AC1 and RM RIm19. The grey shaded area represents the 80^th^ prediction interval derived from 500 simulations the standard model with the RM-specific estimated parameter values. The solid blue line represents the fitted curve. Finally, the blue points are viral load measurements (circle for detectable and crosses for undetectable).

A) **B)**

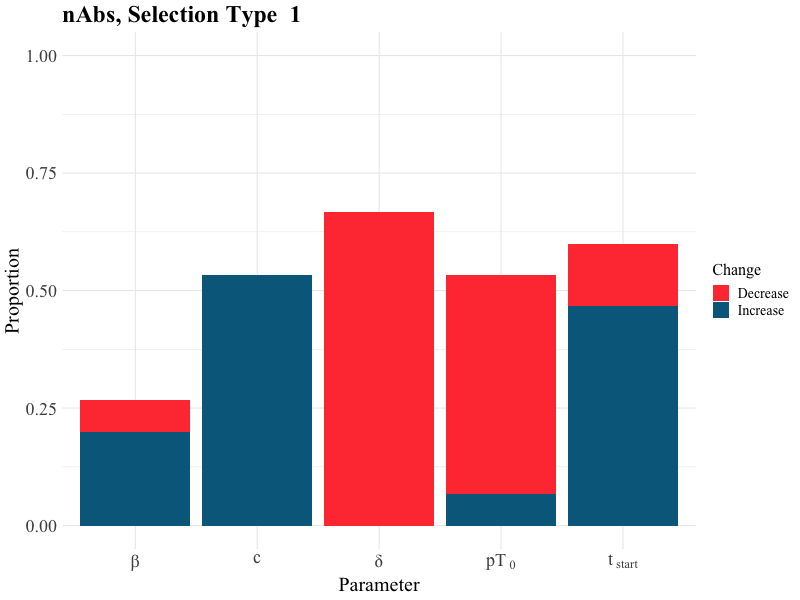

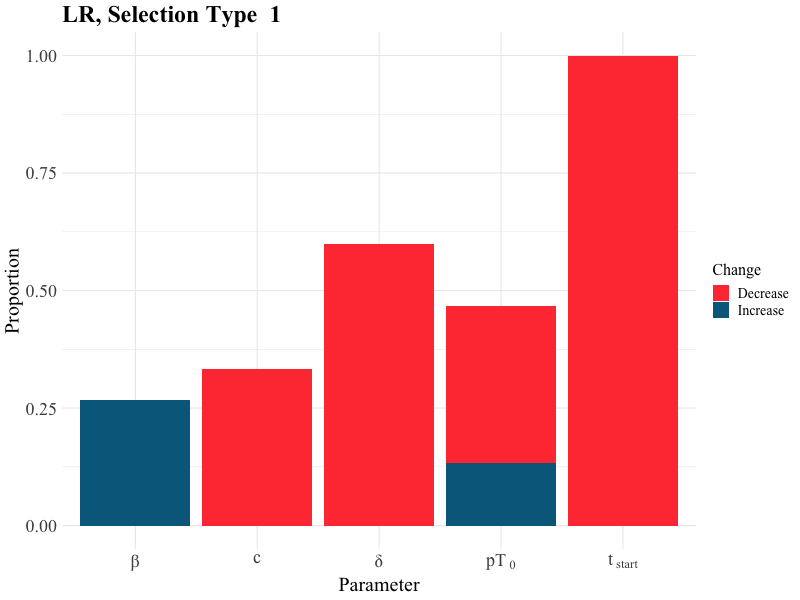

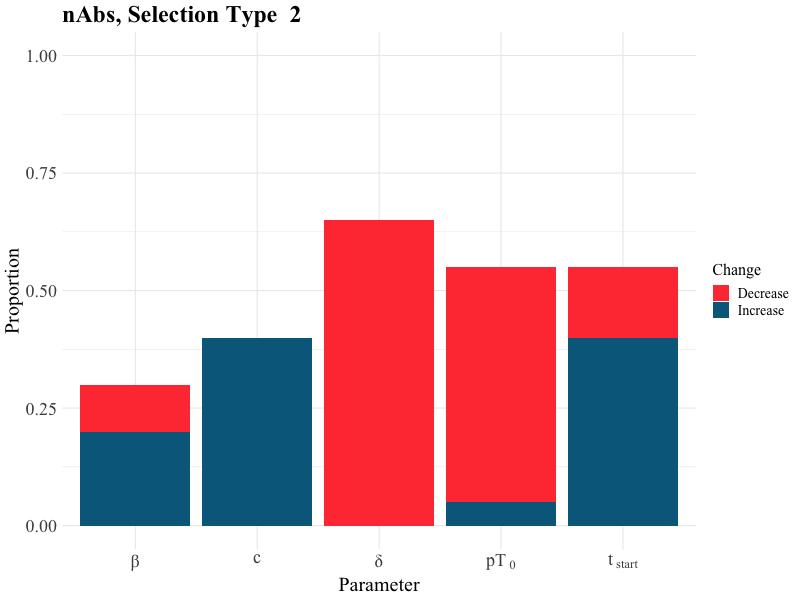

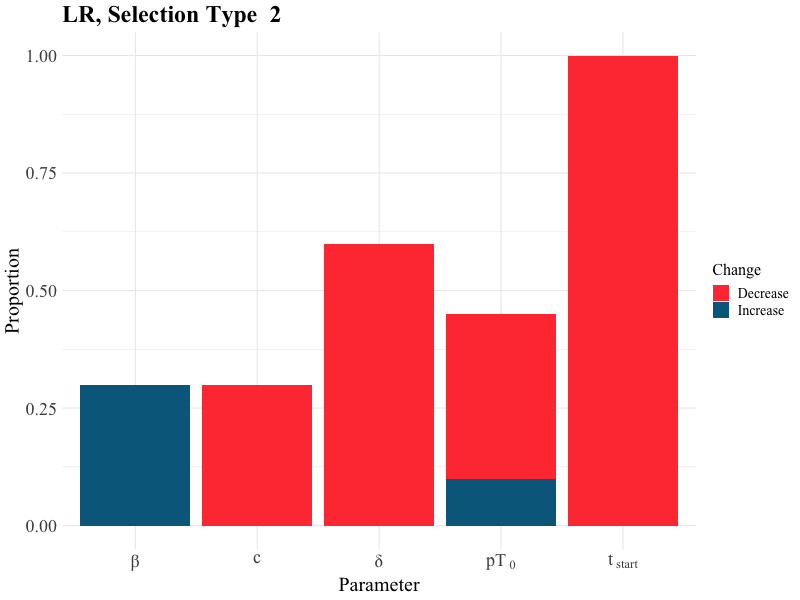

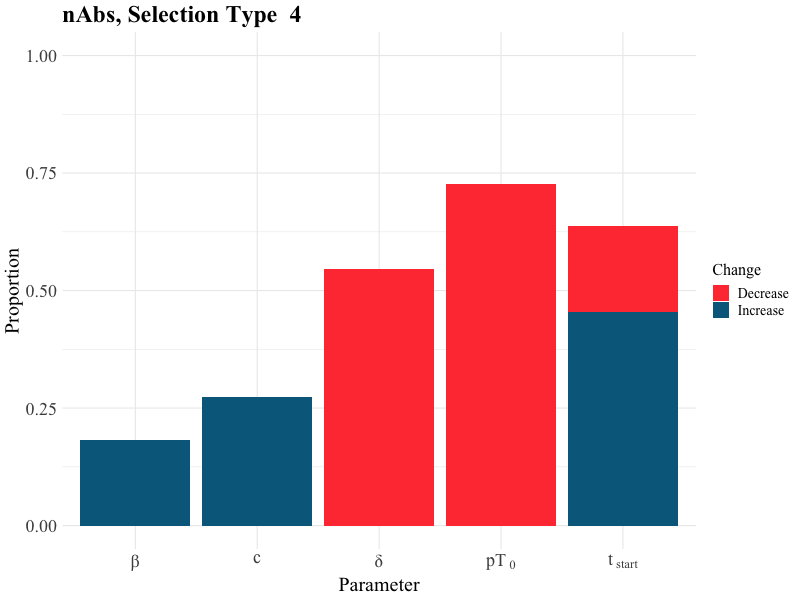

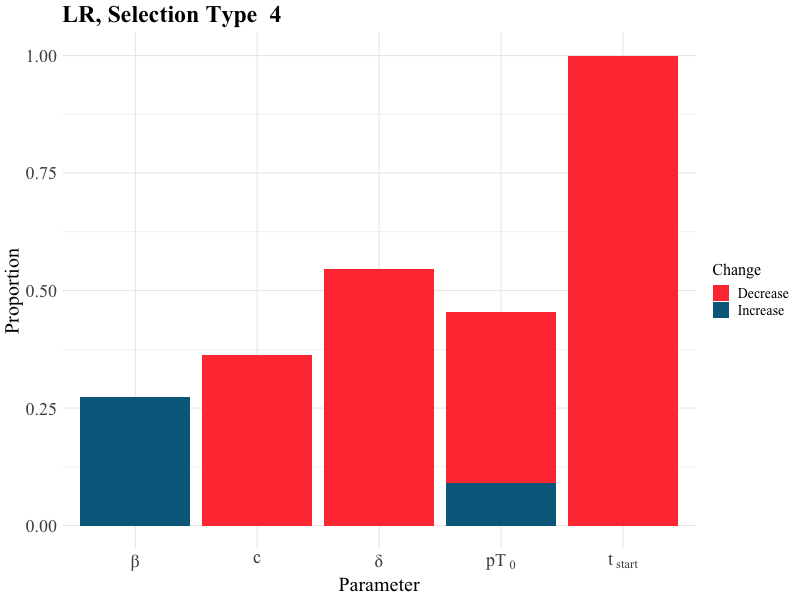

**Supplementary Fig 3**: Barplots depicting the proportion of selected models a potent neutralizing antibody response (nAbs) (A) and detectable latent reservoir (LR) (B) is predicted to affect the median value a model parameter. We also report population-level changes (blue—increase, red—decrease) in the death rate of infected cells (δ), the product of viral production rate and initial number of CD4+ T cells (pT_0_), mass action infectivity (β), and the initiation of exponential viral growth (t_start_). For model selection, we do not focus solely on AIC, but also on additional metrics that we devise, namely the *QPC* and *S*. We evaluate 5 different combinations and thresholds of these three metrics for model selection. For Type 1, we select models with ΔAIC<8, QPC<15 and S<250, obtaining n=15 models. Type 2 includes models with ΔAIC<10, QPC<15 and S<250 (n=20). Finally, type 4 includes models with QPC<14.5 and S<250 (n=11). We also test two types of model selection that neglect QPC. Overall, we find that the associations of the latent reservoir size and parameter changes are consistent among types of selections compared to the effects of neutralizing antibodies. In addition, neglecting the predictive ability of a model, measured through QPC (Types 3 and 5) gives different results compared to types that include QPC.

**A)	B)**

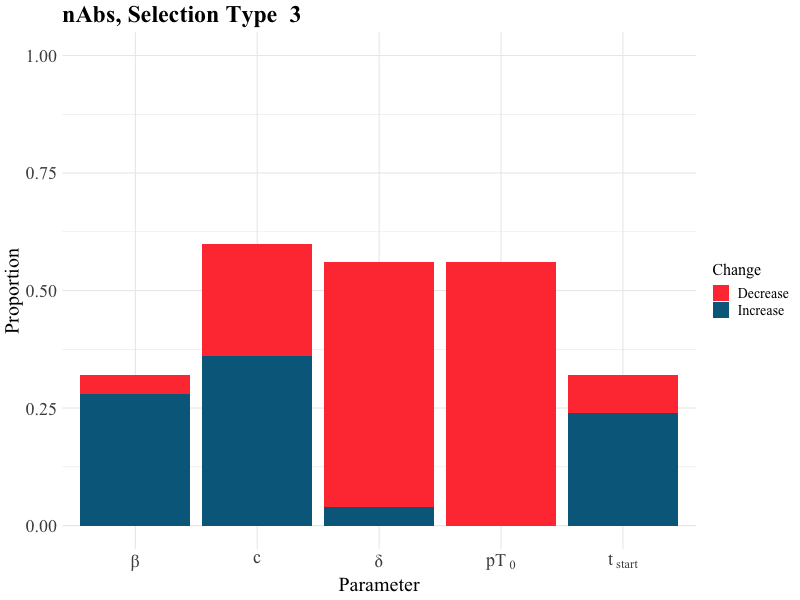

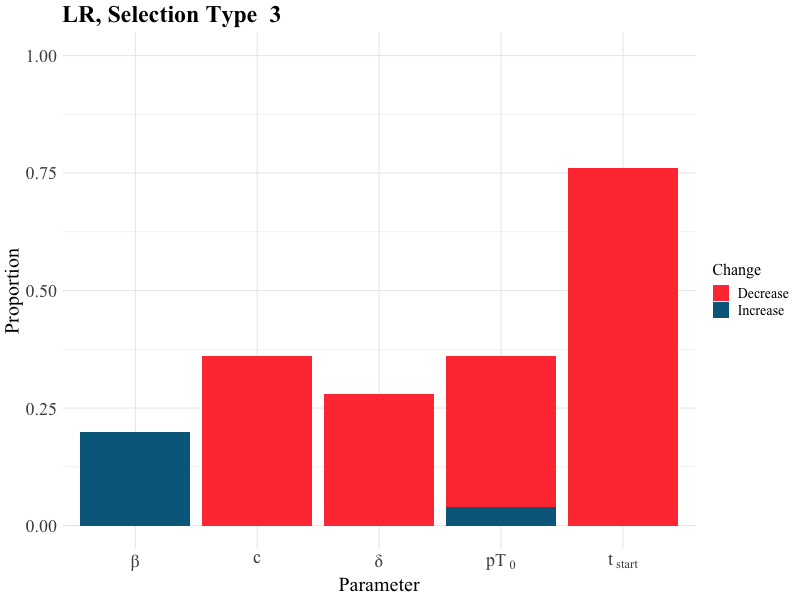

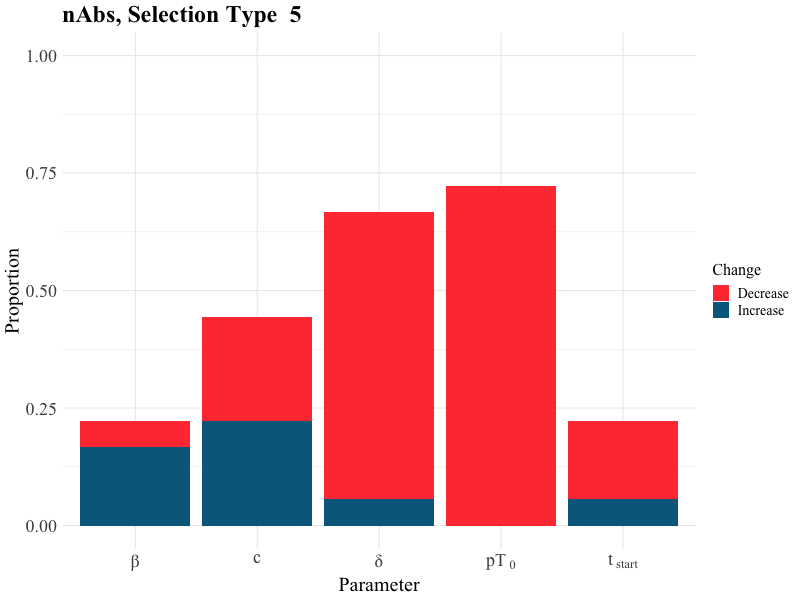

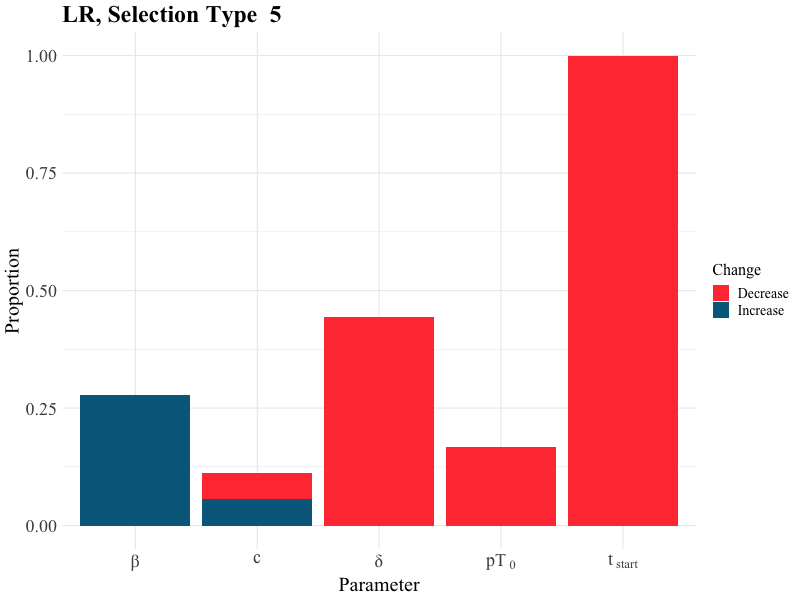

**Supplementary Fig 4**: Barplots depicting the proportion of selected models a potent neutralizing antibody response (nAbs) (A) and detectable latent reservoir (LR) (B) is predicted to affect the median value a model parameter for model selection types that neglect QPC. Type 3 does not impose restrictions on QPC and allows for ΔAIC<8, and S<150, obtaining n=25 models. Finally type 5 focuses on AIC only with selecting models with ΔΑΙC<4 (n=18). We also report population-level changes (blue—increase, red—decrease) in the death rate of infected cells (δ), the product of viral production rate and initial number of CD4+ T cells (pT_0_), mass action infectivity (β), and the initiation of exponential viral growth (t_start_).

**A)**

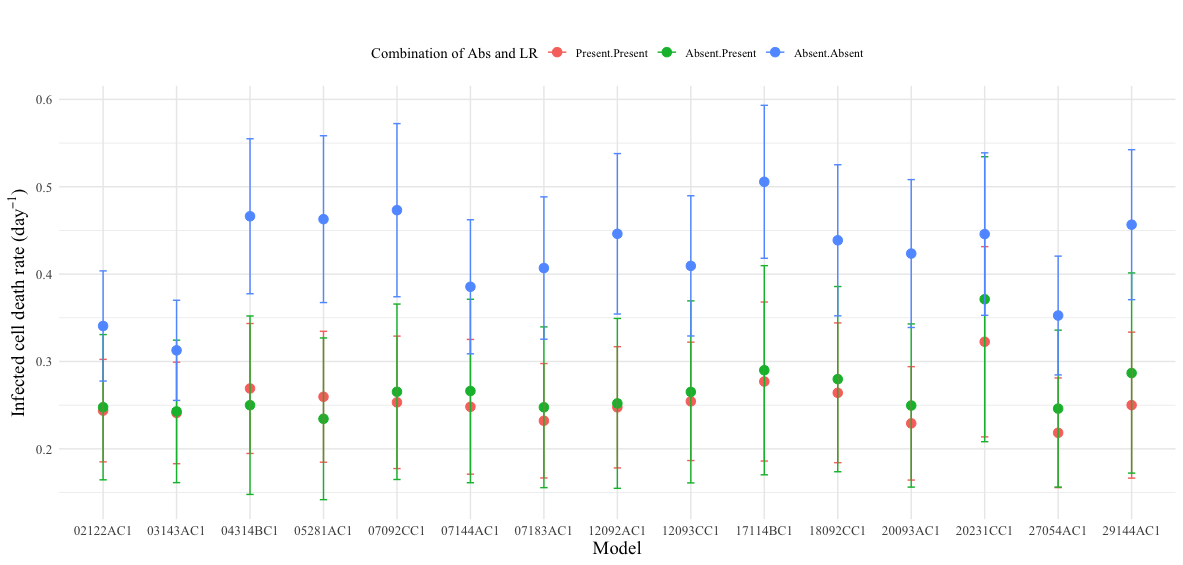

**B)**

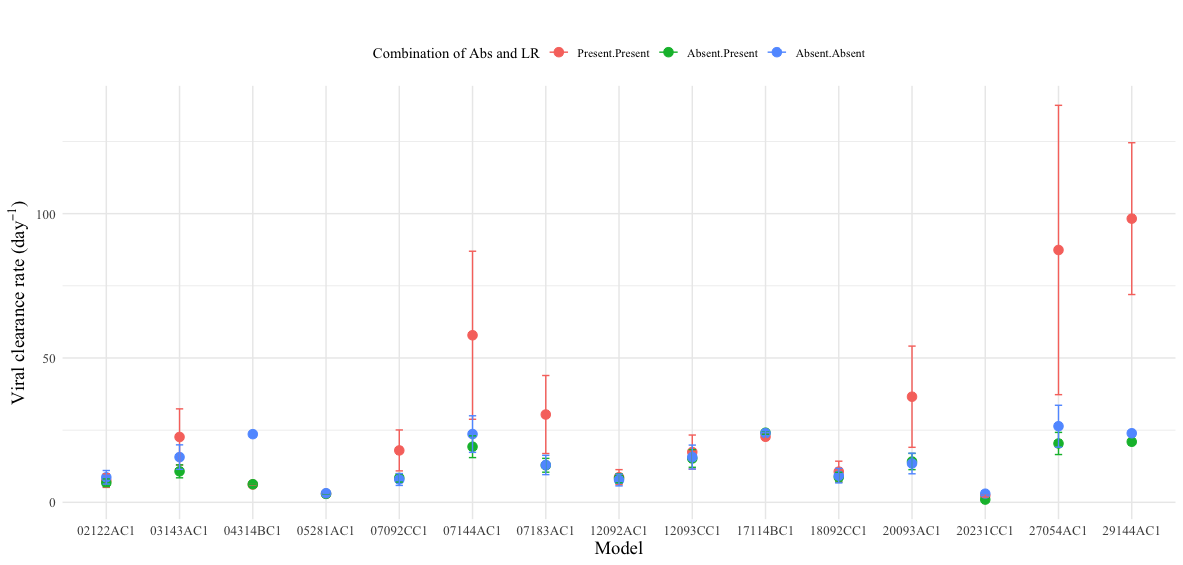

**C)**

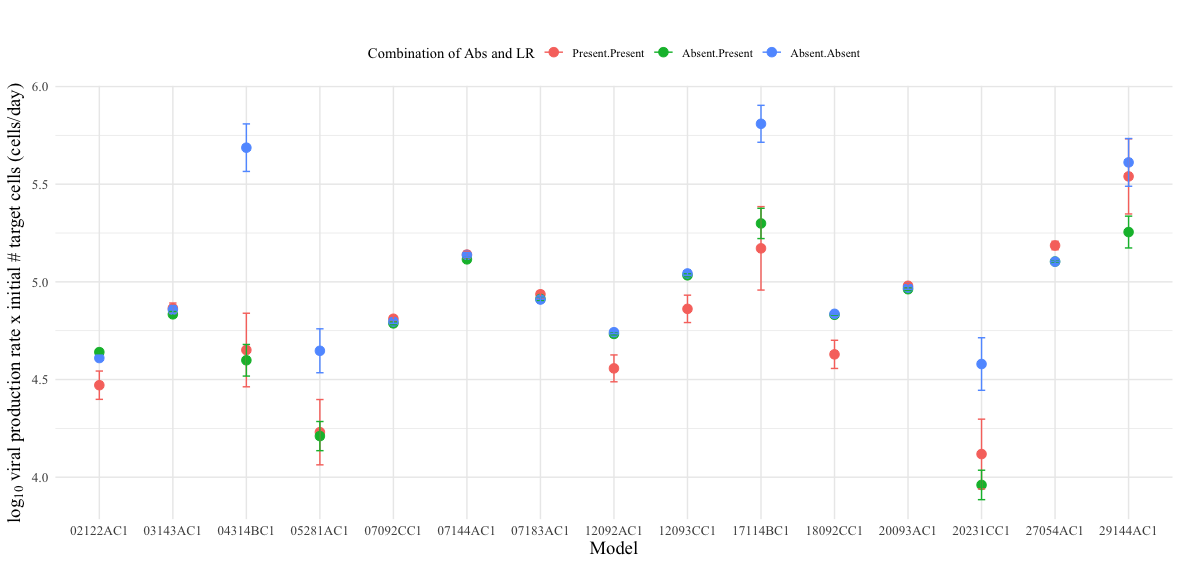

**D)**

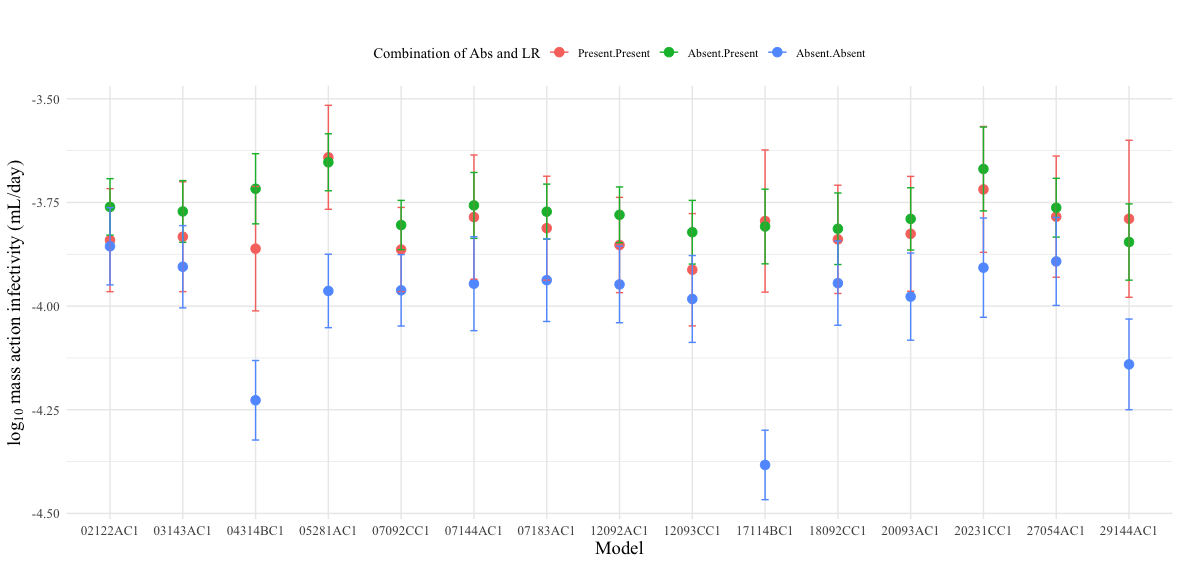

**E)**

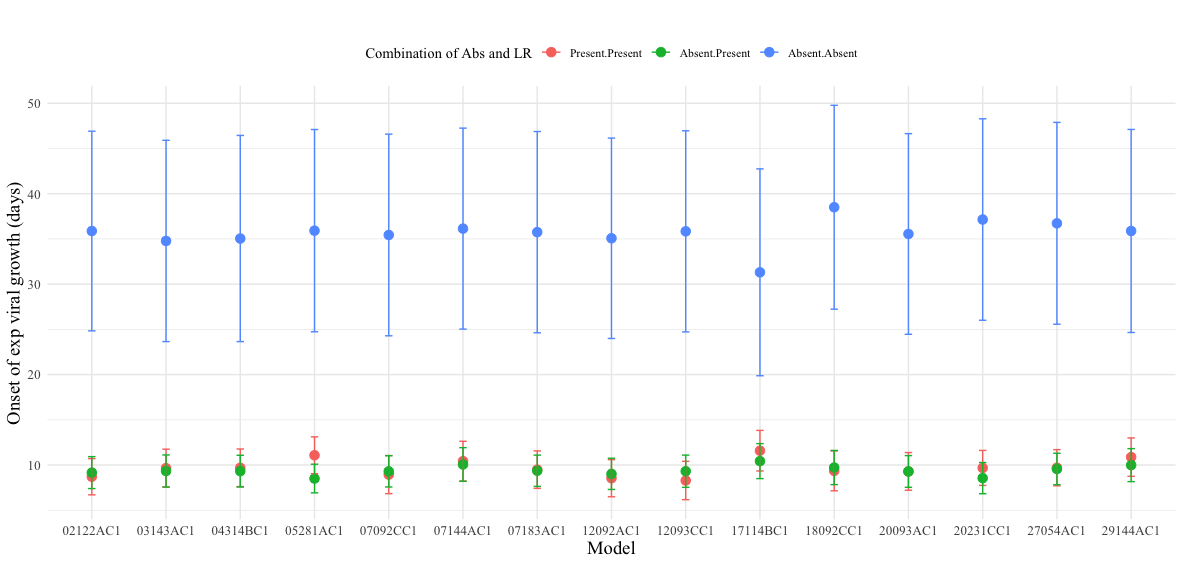

**F)**

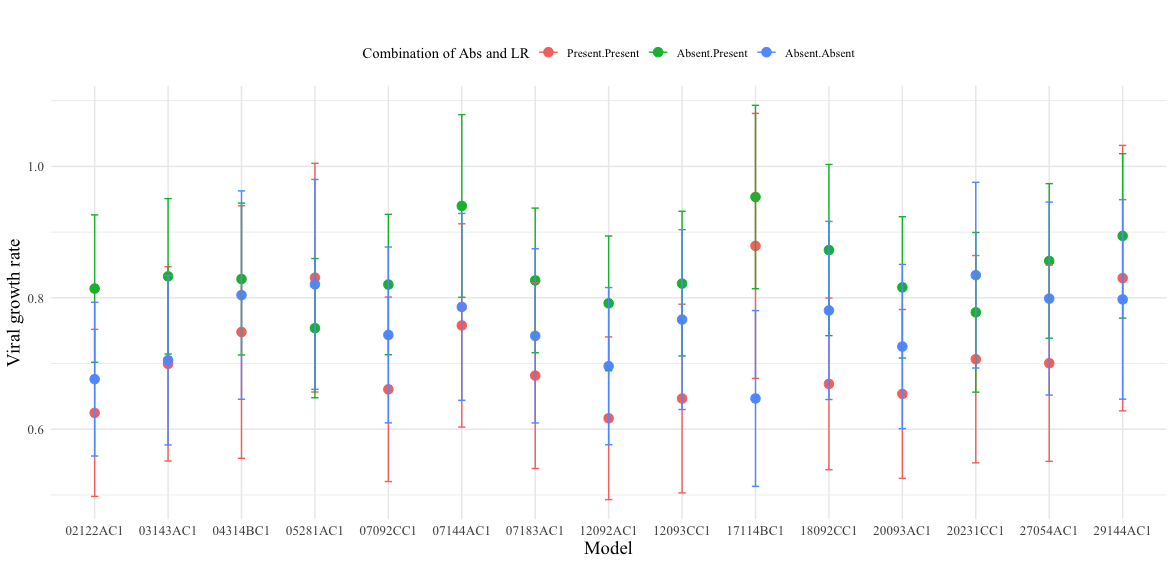

**Supplementary Fig 5:** Mean and variance death rate of infected cells δ (Α), viral clearance rate c (B), the product of the viral production rate and initial number of target cells pT_0_ (C), the mass action infectivity β (D), onset of exponential viral growth t_start_ (E) and viral growth rate (F) for each selected model, grouped by the presence and absence of neutralizing and latent reservoir (red—both nAbs and LR present, blue— both nAbs and LR absent, green—nAbs absent but LR present).

**A)**

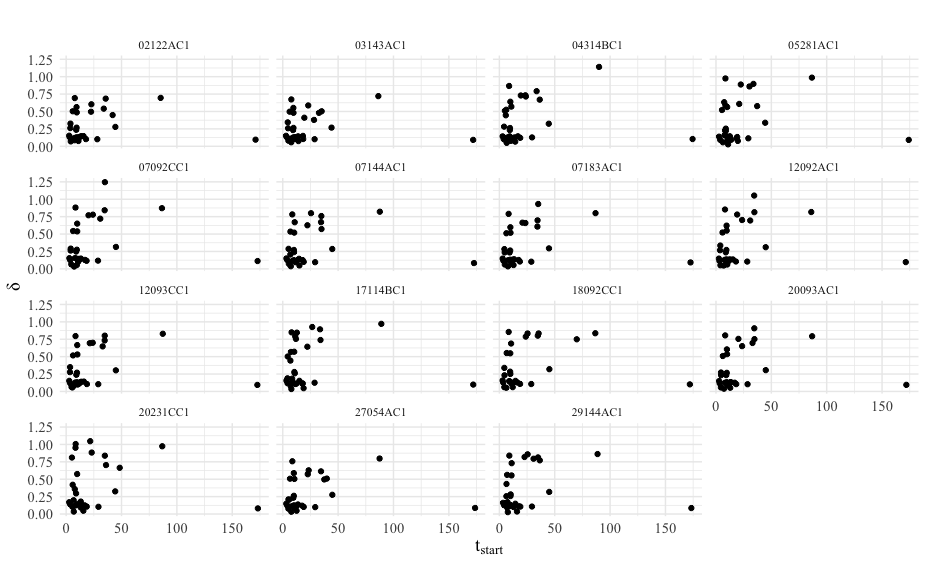

**B)**

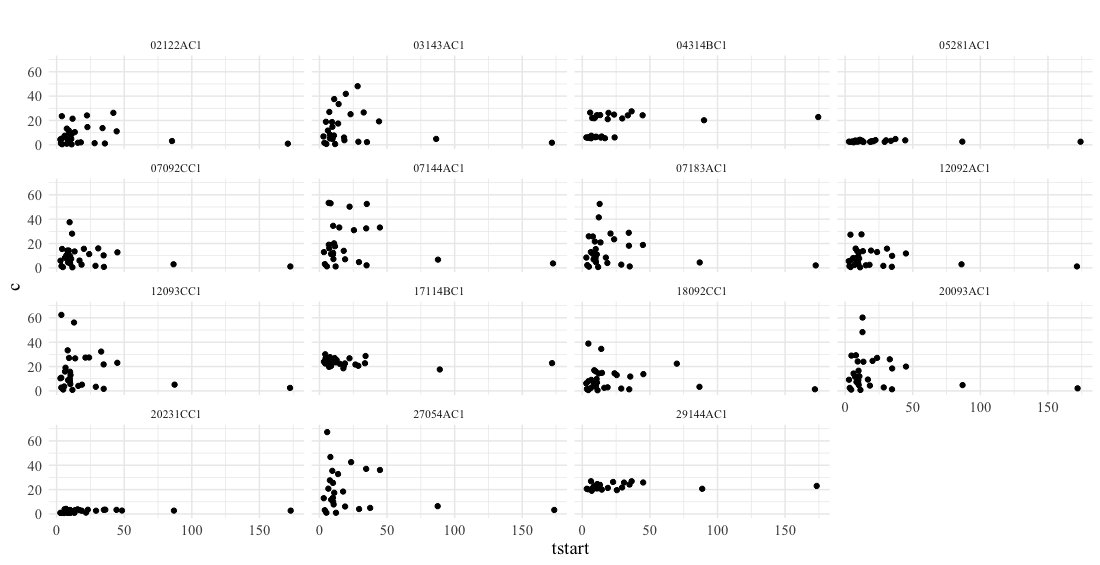

**C)**

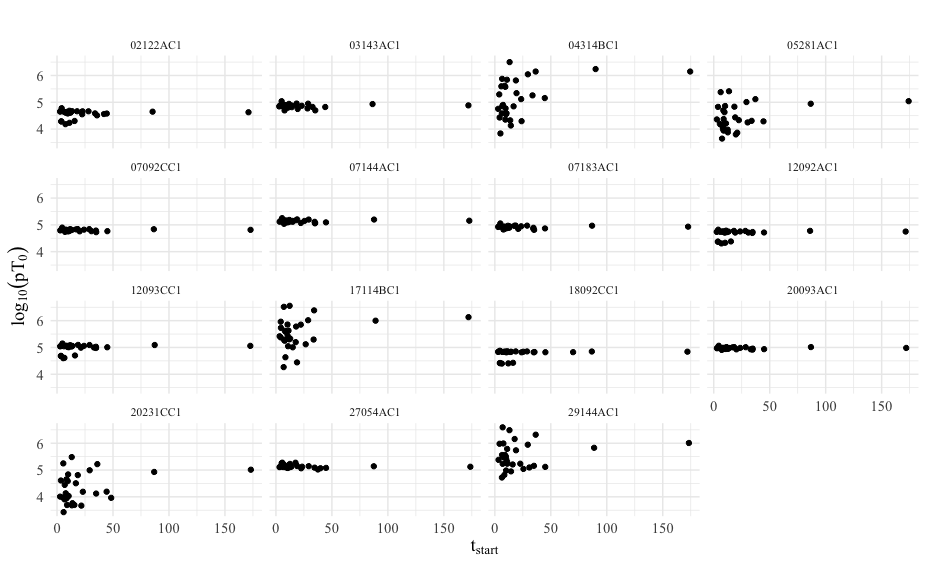

**D)**

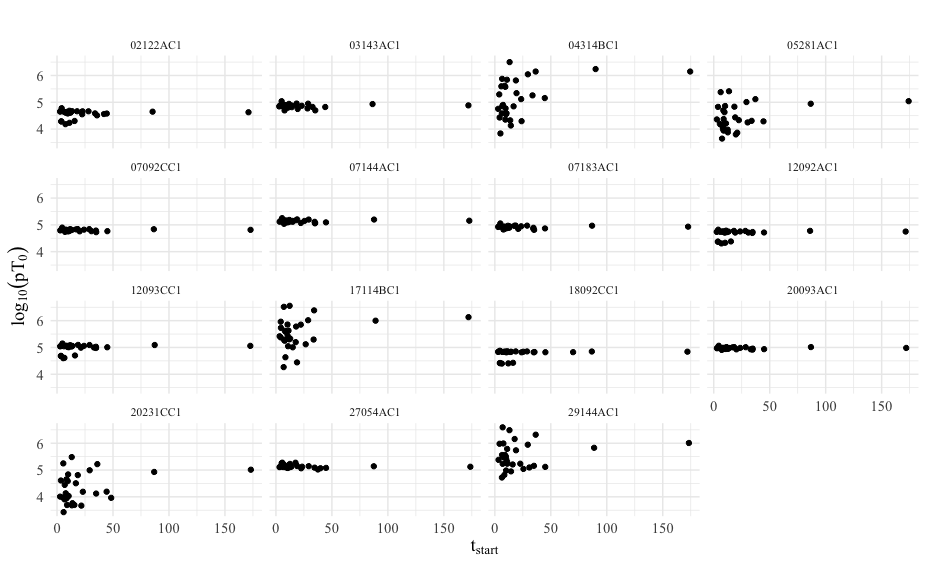

**E)**

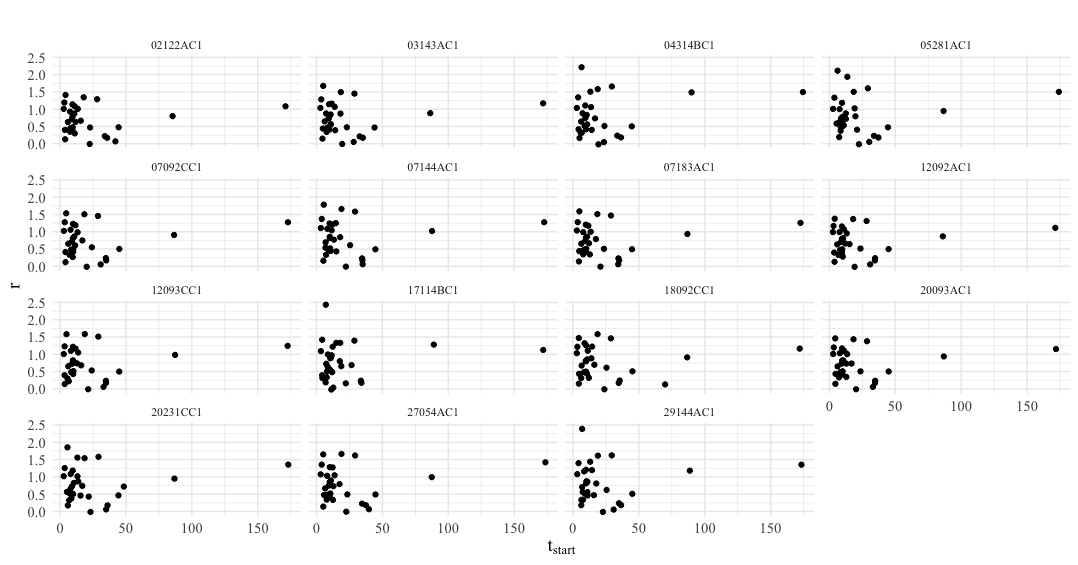

**Supplementary Fig 6:** Scatterplot of the onset of exponential viral growth (t_start_) and death rate of infected cells δ (Α), viral clearance rate c (B), the product of the viral production rate and initial number of target cells pT_0_ (C), the mass action infectivity β (D) and the estimated growth rate of the virus (E) for each selected model.
